# The *Bacillus* Virulome in Endophthalmitis

**DOI:** 10.1101/2020.07.02.184630

**Authors:** Phillip S. Coburn, Frederick C. Miller, Morgan A. Enty, Craig Land, Austin L. LaGrow, Md Huzzatul Mursalin, Michelle C. Callegan

## Abstract

*Bacillus cereus* is recognized as a causative agent of gastrointestinal syndromes, but can also cause a devastating form of intraocular infection known as endophthalmitis. We have previously reported that the PlcR/PapR master virulence factor regulator system regulates intraocular virulence, and that the S-layer protein (SlpA) contributes to the severity of *B. cereus* endophthalmitis. To begin to better understand the role of other *B. cereus* virulence genes in endophthalmitis, expression levels of a subset of factors was measured at the midpoint of disease progression in a murine model of experimental endophthalmitis by RNA-Seq. Several cytolytic toxins were expressed at significantly higher levels *in vivo* than in BHI. The virulence regulators *codY, gntR,* and *nprR* were also expressed *in vivo.* However, at this timepoint, *plcR/papR* was not detectable, we previously reported that a *B. cereus* mutant deficient in PlcR was attenuated in the eye. The motility-related genes *fla, fliF,* and *motB,* and the chemotaxis-related gene *cheA* were detected during infection. We have shown previously that motility and chemotaxis phenotypes are important in *B. cereus* endophthalmitis. The *sodA2* variant of manganese superoxide dismutase was the most highly expression gene *in vivo*, suggesting that this gene is criticial for intraocular survival, potentially through inhibition of neutrophil activity. Expression of the surface layer protein gene, *slpA,* an activator of Toll-like receptors (TLR) −2 and −4, and a potent contributor to intraocular inflammation and disease severvity, was also detected during infection, albeit at low levels. In summary, genes expressed in a mouse model of *Bacillus* endophthalmitis might prove to play crucial roles in the unique virulence of *B. cereus* endophthalmitis, and serve as candidates for novel therapies designed attenuate the severity of this often blinding infection.

**Impact statement:** *B. cereus* causes a potent and rapid infection of the eye that usually results in blindness or enucleation, even with the utilization of current treatment modalities. This necessitates the development of new treatment modalities based on new targets. To begin to better define those *B. cereus* factors with roles in intraocular infection, we analyzed the expression of genes with both known and hypothesized roles in intraocular infection at the midpoint of infection using a murine model of Bacillus endophthalmitis. Potentially targetable candidate genes were demonstrated to be expressed *in vivo*, which suggests that these genes might contribute to the unique virulence of *B. cereus* endophthalmitis. Importantly, our results begin to define the virulome of *B. cereus* in intraocular infections and identify previously uncharacterized factors with potential roles in the severity and outcome of *Bacillus* endophthalmitis.

## Introduction

Endophthalmitis is an infection of the anterior and posterior segments of the eye following the introduction of microorganisms after a surgical procedure (post-operative endophthalmitis [POE]), a traumatic penetrating injury (post-traumatic endophthalmitis [PTE]), or hematogenous spread from an infection of a distant site in the body (endogenous endophthalmitis [EE]) [1–5]. Intraocular infection with *B. cereus* results in devastating inflammation of the eye and, in an unfortunately large number of cases, blindness. *Bacillus* endophthalmitis is one of the most rapidly evolving and severe forms of this infection, and is characterized by fulminant intraocular inflammation and significant vision loss within hours [1–5]. Severe pain, periorbital edema, corneal ring abscesses, anterior chamber inflammation, vitritis, a decreased fundus reflex, propotosis, and a rapid decline in visual acuity are common symptoms of the disease. The destruction of the ocular and retinal architecture has been postulated to result from both bacterial- and host immune-mediated mechanisms. Approximately 70% of patients with *Bacillus* endophthalmitis suffer a significant loss in visual acuity, and half of these require surgical removal of the eye, even with aggressive therapeutic intervention [1–5]. Currently, no universally successful therapeutic regimen has been described to treat this infection. Ineffective antibiotic penetration and contradictory clinical reports on the dose, route, and combination therapy make treating endophthalmitis difficult [1–5]. Moreover, the clinical outcome of this disease is influenced by age, extent of injury, and delays in the initiation of treatment. Examining the mechanisms of both bacterial and host factors in the pathogenesis of *B. cereus* endophthalmitis is of utmost importance in order to develop better therapeutic strategies to treat this blinding disease.

*B. cereus* toxins and cell wall components cause damage to the nonregenerative tissues of the eye during endophthalmitis, either through direct effects on retinal cells or indirectly due to bystander damage from immune cell responses [6–10]. We demonstrated that the PlcR/PapR system, the major transcriptional regulatory system of toxins and virulence in *B. cereus,* contributes to intraocular virulence [9]. However, we could not ascribe individual roles for hemolysin BL, phosphatidylcholine-specific phospholipase C (PC-PLC), and phosphatidylinositol-specific phospholipase C (PI-PLC) in a rabbit model of endophthalmitis [7, 11]. The role of other toxins elaborated by *B. cereus,* including cereolysin O, cytotoxin K, and nonhemolytic enterotoxin (Nhe), have not yet been investigated.

The cell wall of *B. cereus* was more inflammogenic than the cell walls of the intraocular pathogens *S. aureus* and *E. faecalis* in a rabbit model of endophthalmitis [12]. Based on these results, we hypothesized that *B. cereus* cell walls possess a unique structural feature and/or a common component that has evolved structural differences from other Gram-positive pathogens.

A thick peptidoglycan layer, capsular polysaccharide, lipotechoic and teichoic acids, lipoproteins, pili, a glycoprotein S-layer, and flagella comprise the *B. cereus* cell wall [13–15]. *B. cereus* pili interfered with bacterial clearance in a mouse model of endophthalmitis [16], suggesting a role as an antiphagoctyic factor. In the same model, a mutant strain of *Bacillus* deficient in the S-layer protein, SlpA, was less virulent relative to the wild type, parental strain [17]. Moreover, *Bacillus* SlpA stimulated nuclear factor kappa-light-chain-enhancer of activated B cells (NF-κB) in human retinal Muller cells *in vitro,* interfered with phagocytosis by neutrophils and retinal cells, and served as an adhesin to retinal cells [18]. *Bacillus* flagella contribute to the intraocular virulence of *B. cereus* by facilitating migration throughout the eye [12]. However, mutants defective in motility resulted in only a delay in disease progression [8, 19].

We previously reported that *B. cereus* known and putative virulence determinants, virulence-related transcriptional regulators, and superoxide dismutase were expressed in *ex vivo* rabbit vitreous and suggested that these genes might contribute to the unique virulence of *B. cereus* in the eye. In the current study, expression levels of factors known or hypothesized to be involved in virulence and transcriptional regulation, motility, and chemotaxis were measured at the midpoint of infection progression (8 hours postinfection) in an *in vivo* murine model of *B. cereus* endophthalmitis. We also evaluated the expression of these genes after growth in Brain Heart Infusion (BHI) broth at the same time point. The aim of this study was to identify factors expressed *in vivo* that might contribute to pathogenesis and disease progression, and potentially serve as targetable candidates for novel therapeutics. At 8 hours postinfection, significant bacterial growth, inflammation, and retinal function loss are observed in this model. We observed that cytolytic enterotoxins, hemolysins, immune inhibitor metalloproteases, and superoxide dismutases were among the most highly expressed genes in the eye at this time point. These results demonstrated the expression of both putative and known *B. cereus* virulence genes during the midpoint of experimental murine endophthalmitis, and suggested that evaluation of the dynamic virulome during endophthalmitis progression might identify possible targetable candidate genes for novel therapies that aim to reduce virulence.

## Methods

### Bacterial strain and analysis of growth in BHI

Vegetative *B. cereus* ATCC 14579 was cultivated in Luria Burtani (LB) for 18 h at 37°C. The culture was centrifuged for 10 minutes at 4,300 x g, and the bacterial pellet washed 3 times with sterile phosphate-buffered saline (PBS pH 7.4) to remove all traces of LB. After the third wash, the bacterial pellet was resuspended in an equal volume of PBS as the original culture volume and diluted to 200 CFU/μl in PBS prior to intravitreal injection. To assess growth in BHI, bacteria in PBS were diluted to 10^3^ CFU/ml in freshly prepared BHI, and at 8 hours following back-dilution, 20 μl aliquots were diluted 10-fold in sterile PBS. Aliquots from each dilution were plated onto BHI agar plates for bacterial quantification. Bacterial concentrations in CFU/ml were determined and multiplied by 0.4 ml to determine the bacterial concentration in 400 μl of BHI, the volume of PBS used for homogenization of *B. cereus*-infected eyes. Two separate growth analysis experiments were performed using independent batches of BHI, and each experiment was performed with 3 independent cultures. The mean ± the standard error of the mean of the 6 independent cultures is shown.

### Murine endophthalmitis model

This study was conducted in accordance with the recommendations in the Guide for the Care and Use of Laboratory Animals of the National Institutes of Health. The animal use protocol was approved by the Institutional Animal Care and Use Committee of the University of Oklahoma Health Sciences Center (protocol number 18-043). Six week old C57BL/6J mice were acquired from the Jackson Laboratory (Catalog 000664, Bar Harbor ME). Mice were acclimated to conventional housing one week prior to injection to allow for physiological and nutritional stabilization and to equilibrate their microbiota. All mice were housed in microisolation conditions on a 12 h on/12 h off light cycle prior to the experiments under biosafety level 2 conditions during experiments. Mice were 8—10 weeks of age at the time of the experiments.

Mice were anesthetized with a combination of ketamine (85 mg/kg body weight; Ketathesia, Henry Schein Animal Health, Dublin, OH) and xylazine (14 mg/kg body weight; AnaSed; Akorn Inc., Decatur, IL). Intravitreal injections were performed with sterile borosilicate glass micropipettes (Kimble Glass Inc, Vineland, NJ, USA) beveled to an approximate bore size of 10 to 20 μm (BV-10 KT Brown Type micropipette beveller, Sutter Instrument Co., Novato, CA, USA). Eyes were visualized with a stereomicroscope and the micropipettes were inserted just posterior to the superior limbus. *B. cereus* ATCC 14579 at a concentration of 100 CFU in 0.5 μl was injected into the right eyes of 25 mice. Injection rates and volumes were monitored using a programmable cell microinjector (Microdata Instruments, Plainfield, NJ, USA). Triplicate independent experiments consisting of 25 mice per experiment were performed.

### *In vivo* bacterial quantitation

For each *in vivo* experiment, at 8 hours postinfection, all mice were euthanized by CO2 asphyxiation. Eyes were removed, and 5 eyes were placed into each of 5 separate tubes containing 400 *μl* of sterile PBS and 1.0 mm sterile glass beads (Biospec Products Inc., Bartlesville, OK), and homogenized for 60 seconds at 5,000 rpm in a Mini-BeadBeater (Biospec Products, Inc., Bartlesville, OK). Aliquots from each tube were serially diluted and plated on BHI agar plates, and incubated at 37°C. After overnight incubation, the CFU per eye for each tube was calculated by determining the total CFU in 400 *μl* and dividing by 5. The value shown represents the mean ± standard error of the mean of the CFU/eye from the 5 tubes and 3 independent experiments.

### RNA preparation and quantitative PCR Analysis

For the *in vitro* BHI experiments, bacterial cultures were centrifuged for 10 minutes at 4,300 x g and the bacterial cell pellet resuspended in the lysis buffer (RLT) from the RNeasy kit (Qiagen, Germantown, MD). Bacterial cells were then homogenized with sterile 0.1 mm glass beads (Biospec Products Inc., Bartlesville, OK) for 60 seconds at 5,000 rpm in a Mini-BeadBeater (Biospec Products Inc., Bartlesville, OK). Total RNA was purified using the RNeasy kit according to the manufacturer’s instructions (Qiagen). Genomic DNA was removed using the TURBO DNA-free kit (ThermoFisher Scientific, Inc., Waltham, Massachusetts), and ribosomal RNA was depleted using the Ribo-Zero rRNA Removal kit for bacterial rRNA (Illumina, San Diego, CA) according to the manufacturers’ instructions. For total bacterial RNA isolation from *B. cereus*-infected eyes, 5 separate tubes, each containing 5 infected eyes in 400 μl of sterile PBS and 1.0 mm sterile glass beads (Biospec Products Inc., Bartlesville, OK), were homogenized for 60 seconds at 5,000 rpm in a Mini-BeadBeater (Biospec Products, Inc., Bartlesville, OK). Homogenates were centrifuged at 500 x g for 5 minutes to pellet ocular debris, and the supernatants were centrifuged for 10 minutes at 4,300 x g to pellet bacteria. Bacterial pellets were then resuspended in RLT buffer from the RNeasy kit (Qiagen, Germantown, MD), and transferred to tubes with sterile 0.1 mm glass beads (Biospec Products Inc., Bartlesville, OK). Bacteria were homogenized for 60 seconds at 5,000 rpm in a Mini-BeadBeater (Biospec Products Inc., Bartlesville, OK), and bacterial RNA was then purified using RNeasy kit (Qiagen, Germantown, MD). Purified total bacterial RNA from each of the 5 tubes was pooled and genomic DNA removed using the TURBO DNA-free kit (ThermoFisher Scientific, Inc., Waltham, Massachusetts). Ribosomal RNA was depleted using the Ribo-Zero rRNA Removal kit for bacterial rRNA (Illumina, San Diego, CA), and depletion was confirmed via quantitative PCR using primers specific to the *B. cereus* 16s ribosomal RNA as previously described [20].

### RNA sequencing

The triplicate enriched RNA samples obtained from the *in vitro* BHI and the *in vivo* experiments was sequenced using an Illumina MiSeq Next Generation Sequencer at the OUHSC Laboratory for Molecular Biology and Cytometry Research. Raw data for each sample was analyzed using the CLC Genomics Workbench software (Qiagen, Redwood City, CA). Raw sequence reads were mapped to the *B. cereus* ATCC 14579 reference genome for identification of the genes of interest in our study expressed after growth in BHI or *in vivo.* Reads not mapped were excluded further analysis. The number of reads per gene was normalized according to the total number of reads in each library and the gene size. The resulting number was expressed as normalized reads per kilobase per million (RPKM). The values represent the mean RPKM ± the standard deviation of the three independent sequencing runs for each condition.

### Statistics

For *in vitro* growth in BHI, the data is the arithmetic mean ± the standard error of the mean of the CFU/400μl values from two independent experiments with 3 biological replicates per experiment. For the *in vivo* growth experiments, the data is the mean ± standard error of the mean of the CFU/eye from the 5 tubes and 3 independent experiments. For RNA-Seq, the data are the arithmetic means ± the standard deviations of the RPKM values of each gene derived from triplicate *in vitro* BHI and *in vivo* samples. Comparative differences between groups were taken to be statistically significant when p < 0.05. The two-tailed, unpaired t-test with Welch’s correction was used to compare the growth analysis and the RPKM values of the genes in each condition. All statistical analyses were performed using GraphPad Prism 8.2.0 (GraphPad Software, Inc., La Jolla CA).

### Data availability

The RNA-Seq data will be deposited in the Sequence Read Archive at NCBI upon acceptance for publication.

## Results

### Growth of *B. cereus* in BHI and *in vivo*

*In vitro* growth of *B. cereus* ATCC 14579 in BHI 8 hours after back-dilution was compared to *in vivo* growth 8 hours postinfection. As shown in Fig. 1, a statistically significant difference was observed in growth between BHI and in the mouse eye (p=0.0038). These results demonstrate that the total CFU in 400 μl of BHI was approximately a half an order of magnitude higher than the CFU in each eye homogenized in 400 μl of PBS.

**Figure 1.**
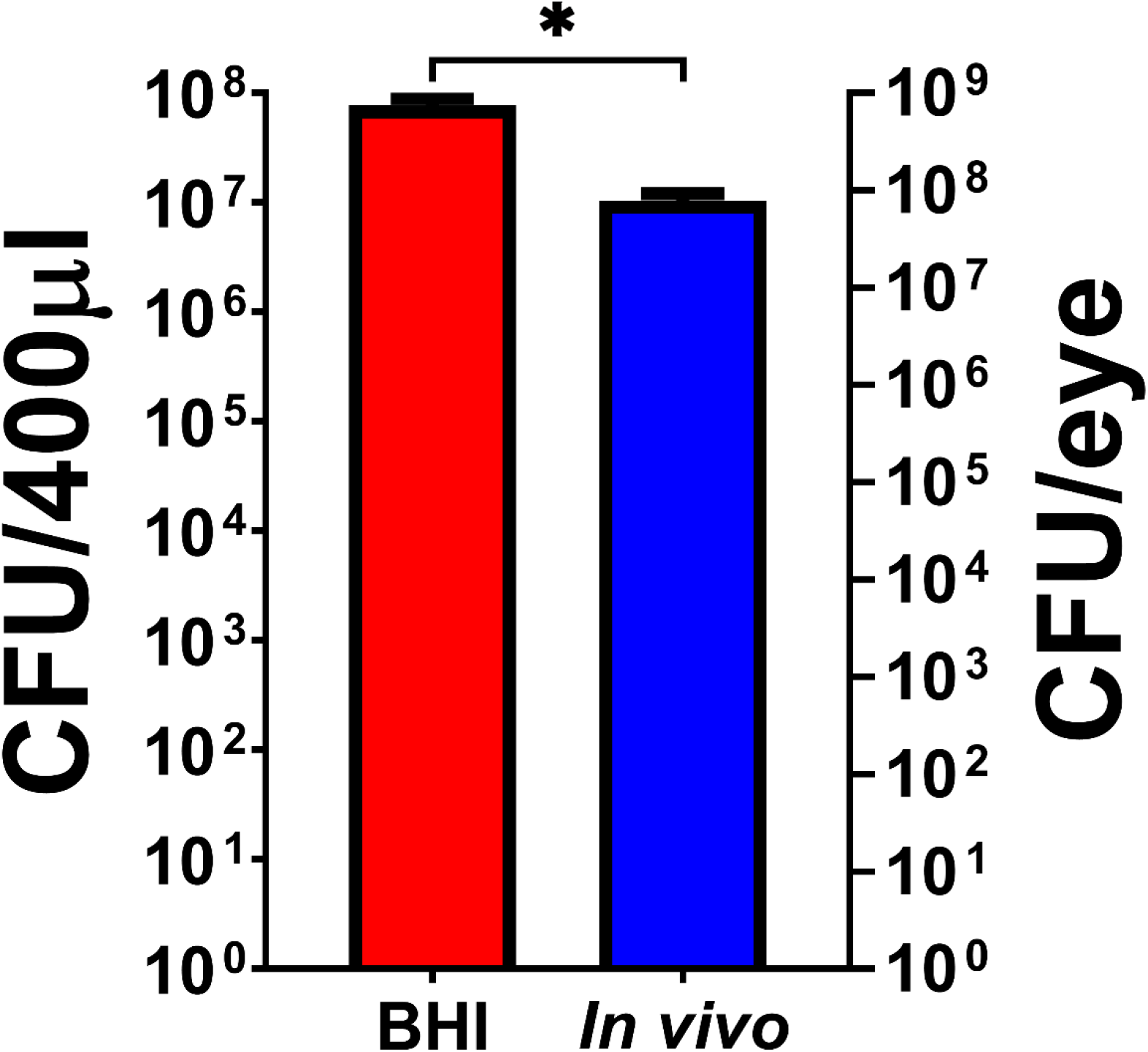
Analysis of the growth of *B. cereus* in BHI and *in vivo.* Growth of *B. cereus* in BHI (Brain Heart Infusion) 8 hours after back dilution, and *in vivo* 8 hours postinfection. The CFU/ml of *B. cereus* 8 hours after back dilution was significantly higher than the concentration in mouse eyes 8 hours after infection (*p=0.0038). The mean CFU/ml ± the standard error of the mean of 6 independent BHI cultures, and the mean CFU/eye ± standard error of the means of 5 tubes (each tube containing 5 eyes) and 3 independent experiments are shown.

### rRNA depletion and RNA-Seq analysis of depleted RNA samples

In the current study, expression levels of factors known and hypothesized to be involved in *B. cereus* endophthalmitis were evaluated during the midpoint of infection in our murine model of endophthalmitis using RNA-Seq. *B. cereus* ATCC 14579 was utilized in this studies since it represents a prototypical strain with a sequenced and annotated genome. The virulence and pathogenesis in endophthalmitis has been previously evaluated and established [9, 16, 21–23]. Similar to our previous results [20], quantitative PCR analysis using primers complementary to the *B. cereus* ATCC 14579 16s rRNA gene demonstrated that the rRNA from the *in vitro* BHI-derived and *in vivo* mouse eye-derived bacterial RNA was depleted (data not shown).

The *B. cereus* ATCC 14579 genome is 5,411,809 bp, and consists of 5,473 genes. The mean read length was 151 bp for all BHI and *in vivo* samples. The mean number of mapped reads and mean percentage of reads that mapped to the *B. cereus* genome for the BHI and *in vivo* samples is shown in Table 1. The mean number of mapped reads for the three BHI-derived RNA samples was 25,505,762 with a mean of 96% of the reads mapping to the *B. cereus* 14579 genome. The mean number of mapped reads for the *in vivo*-derived RNA samples was 12,360,200, and a mean of 99% of the reads mapped to the genome.

**Table 1.** Mean number of reads and percentage of reads that mapped to the *B. cereus* ATCC 14579 genome for each environmental condition.

| Environment | Mean Number of Reads | Percentage of Reads Mapped |
| --- | --- | --- |
| BHI | 25,505,762 | 96% |
| *In vivo* | 12,360,200 | 99% |

### Virulence factor expression *in vivo*

In this study, we assessed the *in vivo* expression of a subset of known and hypothesized virulence genes, transcriptional regulators of virulence-related genes, and genes related to motility and chemotaxis that we previously analyzed *in vitro* and in *ex vivo* vitreous [20]. Expression levels of genes encoding the toxin hemolysin BL (Hbl), the nonhemolytic enterotoxin Nhe, the putative enterotoxins EntA and EntC, entertoxin FM, the putative hemolysin A, cereolysin O, and the metalloproteases InhA1, InhA2, InhA3, and camelysin were measured. The transcriptional regulatory systems related to virulence (SinR/SinI, EntD, CodY, GntR, NprR, and PlcR/PapR), motility-related proteins Fla, FliF, and MotB, and chemotaxis-associated proteins CheA, CheR, and CheY were also evaluated. Superoxide dismutase expression levels *in vivo* were also measured, as our previous data suggested an important role for this factor [20]. Finally, the expression of the surface layer protein gene *(slpA)* was evaluated as this protein is an important contributor to activation of TLR-2 and −4 receptors and initiation of the inflammatory response [18, 24]. Principal component analysis of the RPKM values for each of these mRNAs isolated from *B. cereus* after 8 hours of growth in BHI or 8 hours after intravitreal injection was performed. Fig. 2 shows that the three BHI and *in vivo* replicates clustered into distinct gene expression patterns. This suggested reproducible results among the replicates of each condition.

**Figure 2.**
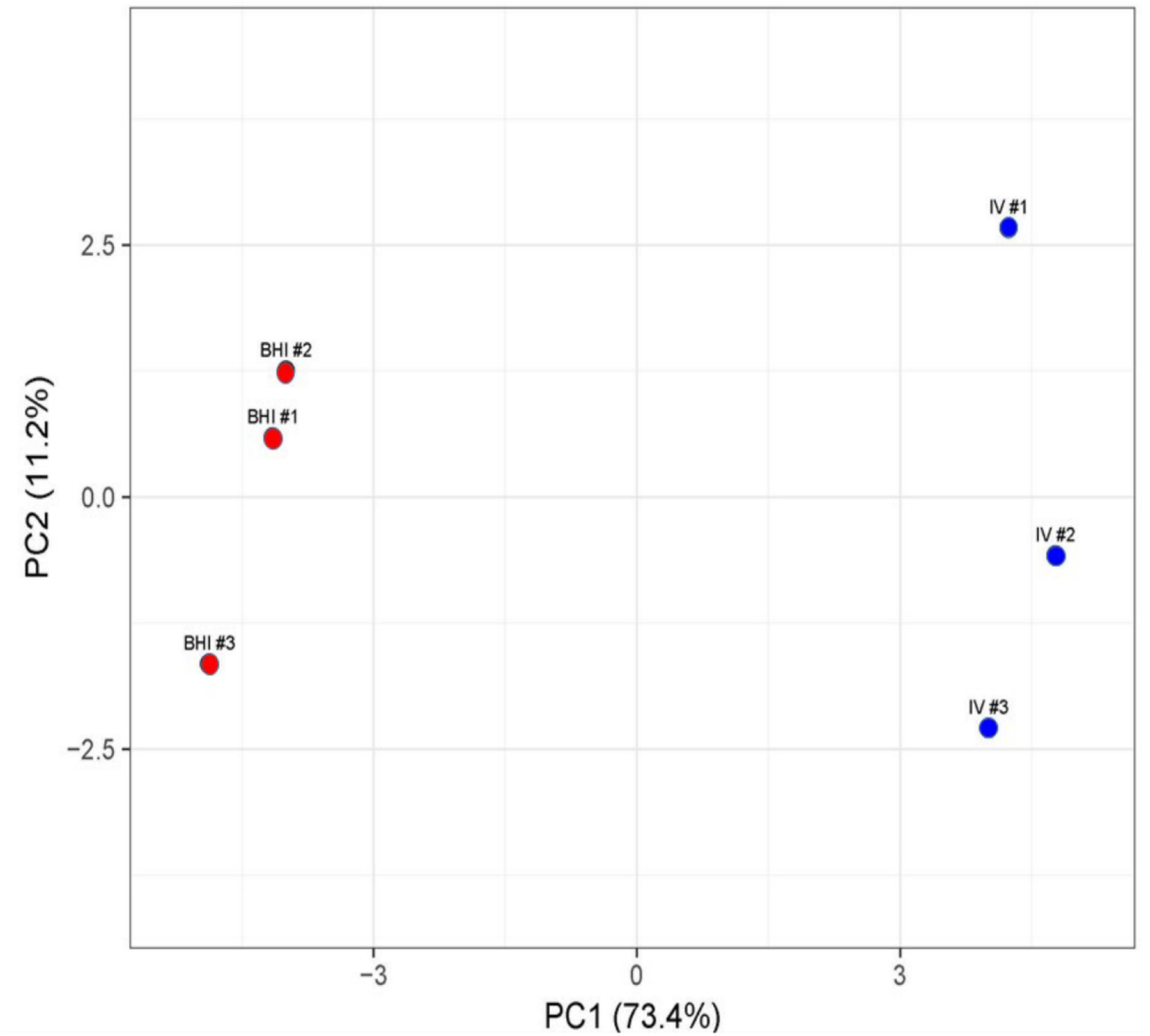
Principal component analysis of *B. cereus* gene expression in BHI or *in vivo.* The expression levels of genes related to virulence, regulation of virulence, chemotaxis, and motility from each of the three independent RNA-Seq runs of *B. cereus* cultivated in BHI 8 hours after back-dilution or 8 hours following intravitreal infection of mouse eyes was analyzed by principal component analysis. Gene expression levels in BHI from each of the 3 independent RNA-Seq experiments clustered distinctly from gene expression levels in infected mouse eyes from each of the 3 independent RNA-Seq experiments.

Toxin expression at 8 hours postinfection was observed at levels similar or higher than in BHI. Among those that were expressed *in vivo* were Hbl, Nhe, EntA, EntC,EntFM HlyA, CerO, InhA1, and InhA2. Mean RPKM values of *hblL1, hblL2,* and *hblB* in vivo were 744, 1002, and 1543, respectively (Fig. 3A). Expression of these genes was 21-, 36-, and 38-fold higher *in vivo* than in BHI (p≤0.0173), respectively. Mean RPKM of the *nheL1* and *nheL2* genes was 1694 and 410, respectively (Fig. 3B). Expression of *nheL1* was 26-fold (p<0.0001) and *nheL2* was 7-fold (p=0.0628) higher *in vivo* than in BHI, with the latter not being significant. Mean RPKM values of 1182 for *entA,* and 1533 for *entC* were observed *in vivo* (Fig. 3C). Expression of *entA* was significantly higher *in vivo* than in BHI (p=0.0068), however *entC* expression was not significantly different *in vivo* than in BHI (p=0.1827). The *entFM* gene was expressed at similar levels *in vivo* (mean RPKM = 1503) and BHI (mean RPKM = 1839) (p=0.6257) (Fig. 3C). The HlyA-encoding gene was expressed at nearly identifical levels *in vivo* (mean RPKM = 364) and BHI (mean RPKM = 379) (p=0.8876) (Fig. 3D). The cholesterol dependent cytolysin (CDC) cereolysin O gene was 9-fold higher *in vivo* than in BHI (p=0.0050) (Fig. 3D).

**Figure 3.**
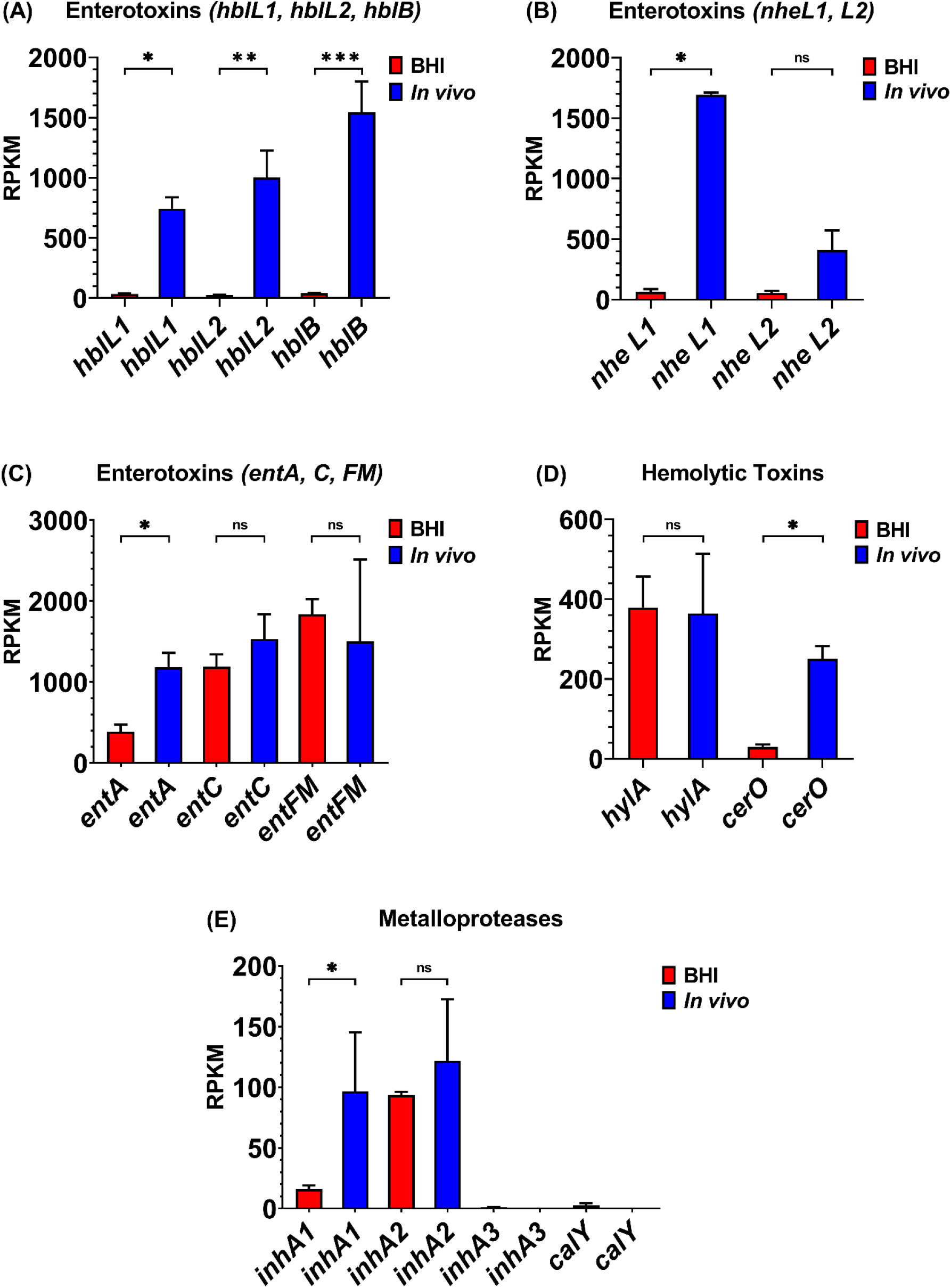
Normalized virulence-related gene expression in BHI or *in vivo.* Reads per kilobase per million (RPKM) for the hemolysin BL genes *hblL1* (*p=0.0057), *hblL2* (**p=0.0173), and *hblB* (***p=0.0096) *in vivo* relative to BHI (A); nonhemolytic enterotoxin genes *nheL1* (*p<0.0001) and *nheL2* (ns = not significant) *in vivo* relative to BHI (B); the enterotoxin genes *entA* (*p=0.0068), *entC* (ns), and the *entFM* (ns) *in vivo* relative to BHI (C); the *hlyA* (ns) and *cerO* (*p=0.0050) genes *in vivo* relative to BHI (D); and the metalloprotease *inhA1* (*p=0.0270), *inhA2* (ns), *inhA3* (not detected), and *calY* (not detected) genes *in vivo* relative to BHI (E). BHI is shown in red, and *in vivo* in blue. RPKM values are the means ± the standard deviations of three independent RNA-Seq runs.

The gene expression of the metalloproteases InhA1, InhA2, InhA3, and camelysin were measured *in vivo* relative to BHI. The *inhA1* and *inhA2* genes were expressed *in vivo* to a similar degree, with a mean RPKM of 97 for *inhA1* and 122 for *inhA2* (Fig. 3E). Transcript levels of *inhA1* were 6-fold higher *in vivo* than in BHI (p=0.0270), however, *inhA2* levels were not significantly different between the environments (p=0.4350). In both environments, expression of *inhA3* was not detected (Fig. 3E), as expected since this strain does not express *inhA3* due to a mutation in *npr* [25]. The *calY* gene, which encodes a cell surface-associated metalloproteinase, camelysin, was also not expressed in either environment (Fig. 3E), although we previously observed expression in LB and *ex vivo* vitreous [20]. These results indicate that with exception of *inhA3* and *calY,* expression of both known and putative *B. cereus* virulence factors occurred *in vivo*, with expression levels being higher than or similar to BHI.

### Virulence-related transcriptional regulatory gene expression *in vivo*

Expression of genes reported to be involved in virulence factor regulation was highly variable in BHI and at 8 hours postinfection *in vivo* (Fig. 4). The biofilm formation- and enterotoxin-related regulatory gene *sinR* was not detected at this time point *in vivo* (Fig. 4A) but was significantly higher in BHI (p= 0.0024). Expression of the SinR inhibitor *sinI* was not detected *in vivo* or in BHI (Fig. 4A). Similarly, expression of *entD* was not detected in both environments (Fig. 4B). In BHI, *codY* levels were 2-fold higher than *in vivo* (p=0.0196) (Fig. 4B). While expression of *gntR* was higher *in vivo* than in BHI, this difference was not significant (p=0.2884, Fig. 4C). However, *nprR* expression was 56-fold higher *in vivo* relative to BHI (p=0.0014, Fig. 4C). Of considerable interest is that the expression of the master toxin and virulence gene regulatory system *plcR/papR* was not detected 8 hours postinfection (Fig. 4D). Mean RPKM values of 10 for *plcR* and 45 for *papR* were observed in BHI (Fig. 4E). The *plcR* gene expression level was not significantly higher in BHI than *in vivo* (p=0.0619); however *papR* expression was significantly higher in BHI than *in vivo* (p=0.0025). In summary, only *codY, gntR,* and *nprR* were expressed at 8 hours postinfection *in vivo,* with the regulators *gntR* and *nprR* being the most highly expressed regulators in the eye.

**Figure 4.**
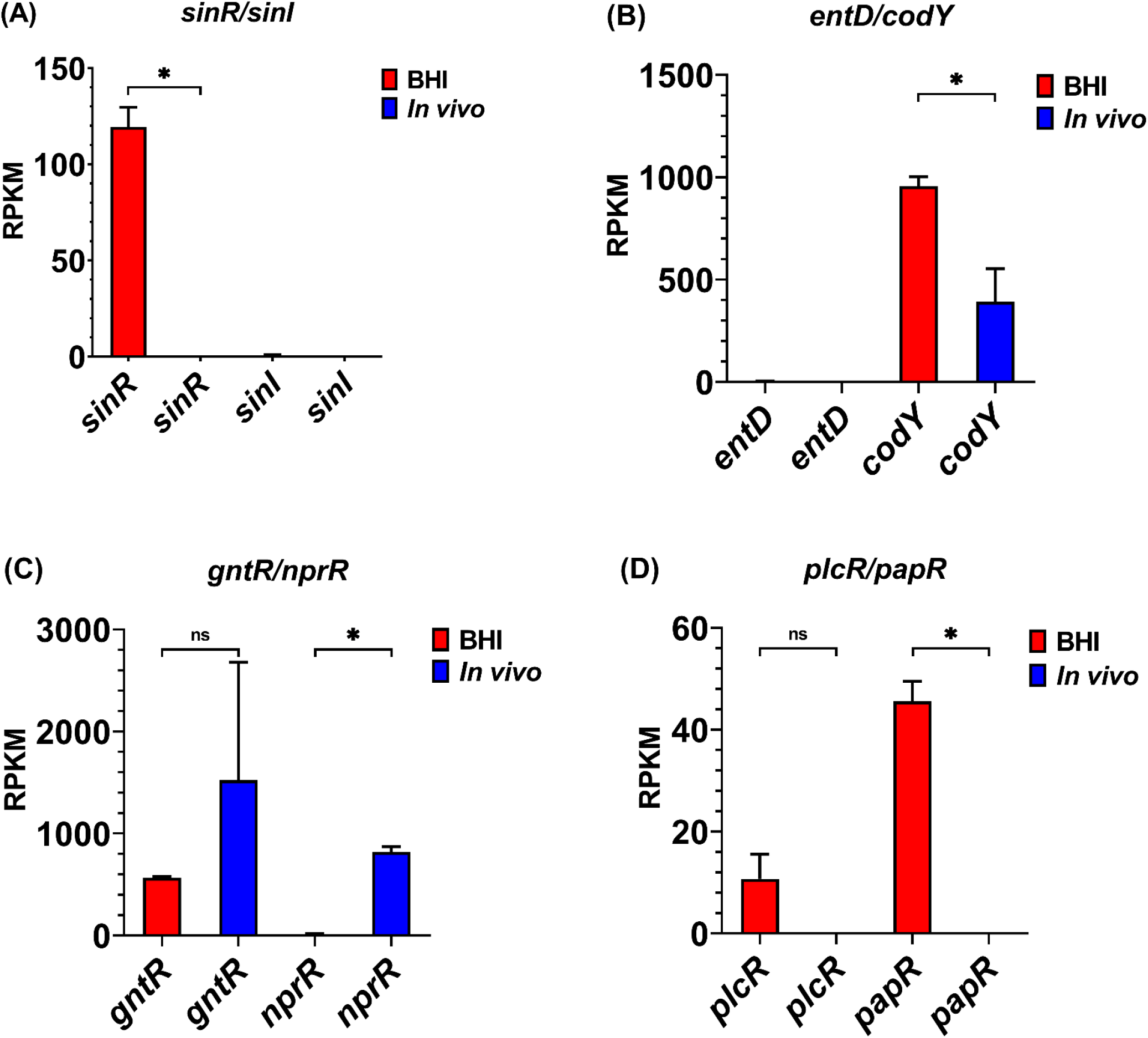
Normalized transcriptional regulatory gene expression in BHI or *in vivo.* Reads per kilobase per million (RPKM) for the transcriptional regulator genes *sinR* (*p= 0.0024) in BHI relative to *in vivo,* and *sinI* (not detected) (A); the genes *entD* (not detected) and *codY* (*p=0.0196) in BHI relative to *in vivo* (B); the genes *gntR* (ns = not significant) and *nprR* (*p=0.0014) *in vivo* relative to BHI (C); and the genes *plcR* (ns) and *papR* (*p=0.0025) in BHI relative to *in vivo* (D). BHI is shown in red, and *in vivo* in blue. RPKM values are the means ± the standard deviations of three independent experiments RNA-Seq runs.

### Expression of genes associated with motility and chemotaxis *in vivo*

*B. cereus* motility contributes to endophthalmitis and requires flagellar activity [8, 19]. The mean RPKM for the flagellar stator protein gene, *motB*, *in vivo* was 166, the flagellar M-ring gene *fliF* was 253, and the flagellin subunit gene *fla* was 785 (Fig. 5A). Expression of *motB* and *fliF* were 2-fold (p≤0.0147), and *fla* was 7-fold (p<0.0001) higher in BHI than *in vivo*. The mean RPKM of *cheA, cheY,* and *cheR* was 139, 0, and 0, respectively, *in vivo* (Fig. 5B). Expression levels of *cheA, cheY,* and *cheR* were significantly higher in BHI (p≤0.0415). Expression levels of the motility-, but not chemotaxis-related genes in the eye correlated with motility and migration of *B. cereus* within the eye at this time point during endophthalmitis.

**Figure 5.**
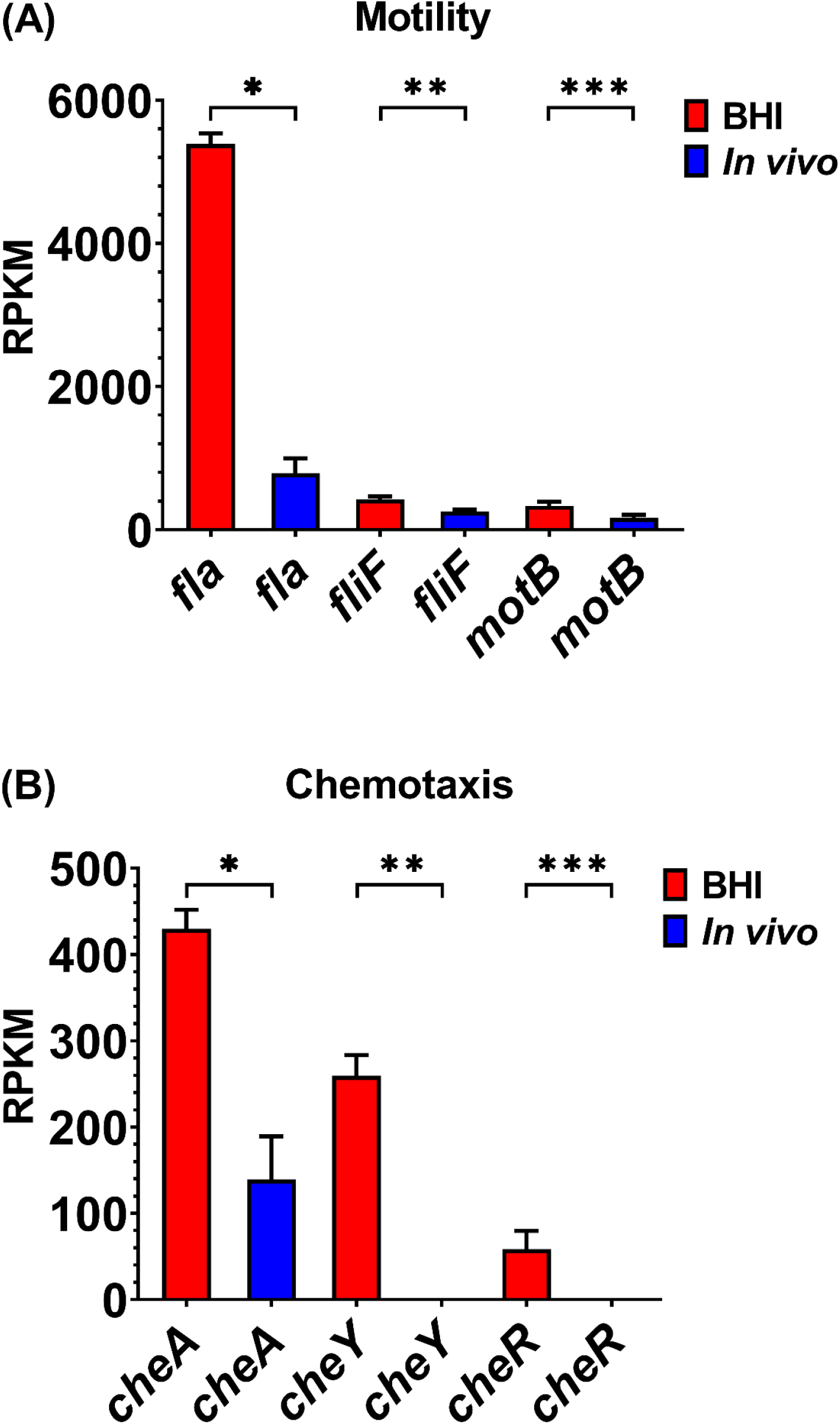
Normalized motility and chemotaxis-related gene expression in BHI or *in vivo.* Reads per kilobase per million (RPKM) for the motility-related genes *fla* (*p<0.0001), *fliF* (**p=0.0064), and *motB* (***p=0.0147) in BHI relative to *in vivo* (A); and the chemotaxis-related genes *cheA* (*p=0.0038), *cheY* (**p=0.029), and *cheR* (***p=0.0415) in BHI relative to *in vivo* (B). BHI is shown in red, and *in vivo* in blue. RPKM values are the means ± the standard deviations of three independent RNA-Seq runs.

### Expression of superoxide dismutase (SodA) *in vivo*

The genes *sodA1* and *sodA2,* encoding two variants of manganese superoxide dismutase, were highly expressed during infection of the eye. The mean RPKM for *sodA1* was 1,891 and for *sodA2* was 16,721 *in vivo* (Fig. 6), the latter being the most highly expressed gene *in vivo.* Expression of *sodA1 in vivo* was not significantly than in BHI (p=0.4608), but expression of *sodA2* was significantly higher *in vivo* than after growth in BHI (p=0.0460). The high level expression of *sodA2* observed at 8 hours postinfection suggested that superoxide dismutase might protect *B. cereus* from internally-produced superoxide anion during growth, or possibly from superoxide generated by neutrophils, the expression of which was not assessed in this study.

**Figure 6.**
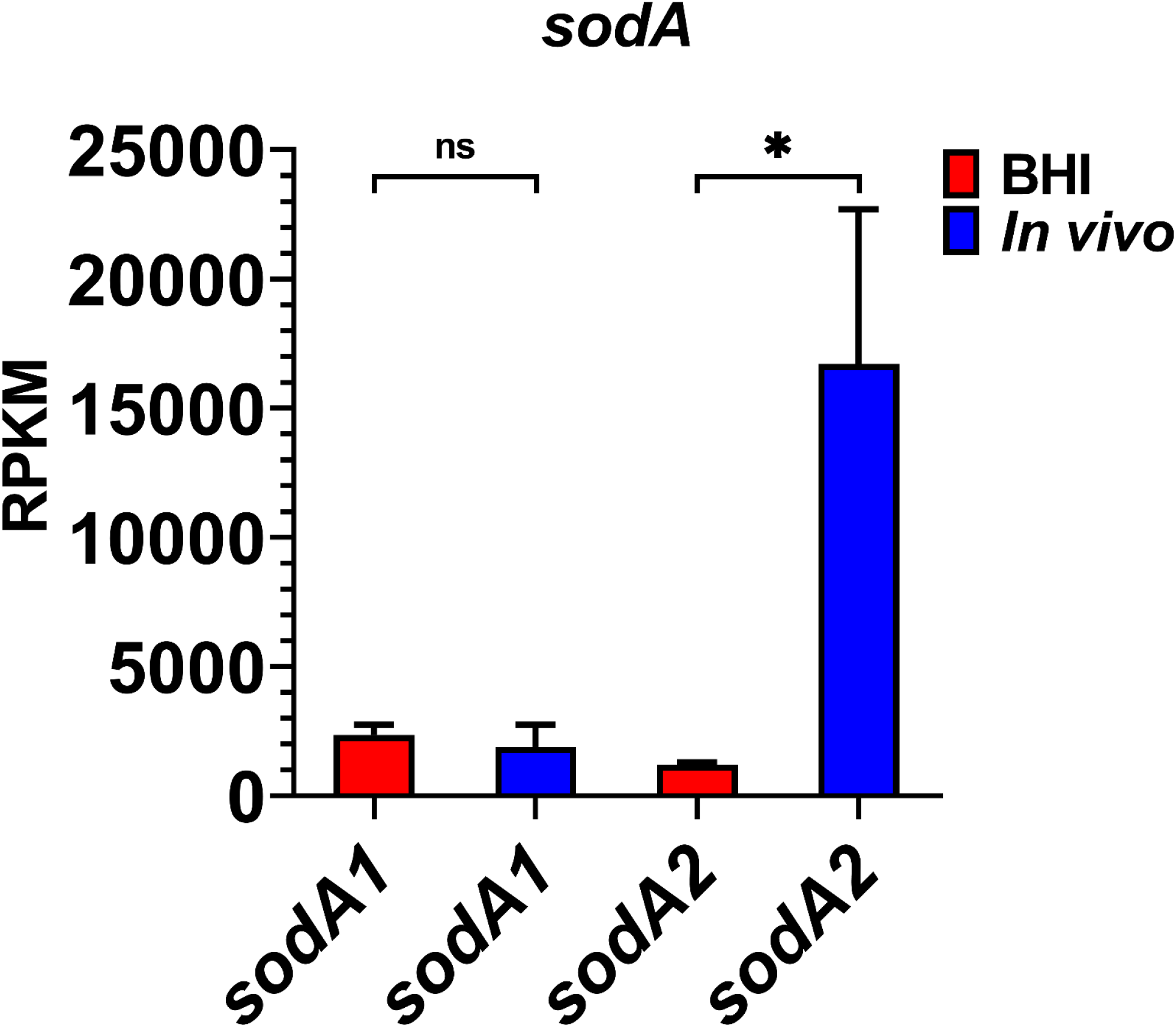
Normalized superoxide dismutase (SodA) gene expression in BHI or *in vivo.* Reads per kilobase per million (RPKM) for the genes *sodA1* (ns = not significant) and *sodA2* (*p=0.0460) *in vivo* relative to BHI. BHI is shown in red, and *in vivo* in blue. RPKM values are the means ± the standard deviations of three independent RNA-Seq runs.

### Expression of surface layer protein (SlpA) *in vivo*

The *Bacillus* surface layer protein, SlpA, has been shown to play a key role in activation of both TLR-2 and −4 and in the severity of infection in a mouse model of *Bacillus* endophthalmitis [17, 18, 24]. The mean RPKM of *slpA* was 25 *in vivo,* and 36 after growth in BHI (Fig. 7). The means were not statistically different (p=0.3777). While this level is relatively low compared to the expression of other known virulence factors *in vivo,* detection of *slpA* transcripts during infection is consistent with our results showing that *slpA* contributes to the inflammatory response and disease severity in a mouse model of *Bacillus* endophthalmitis.

**Figure 7.**
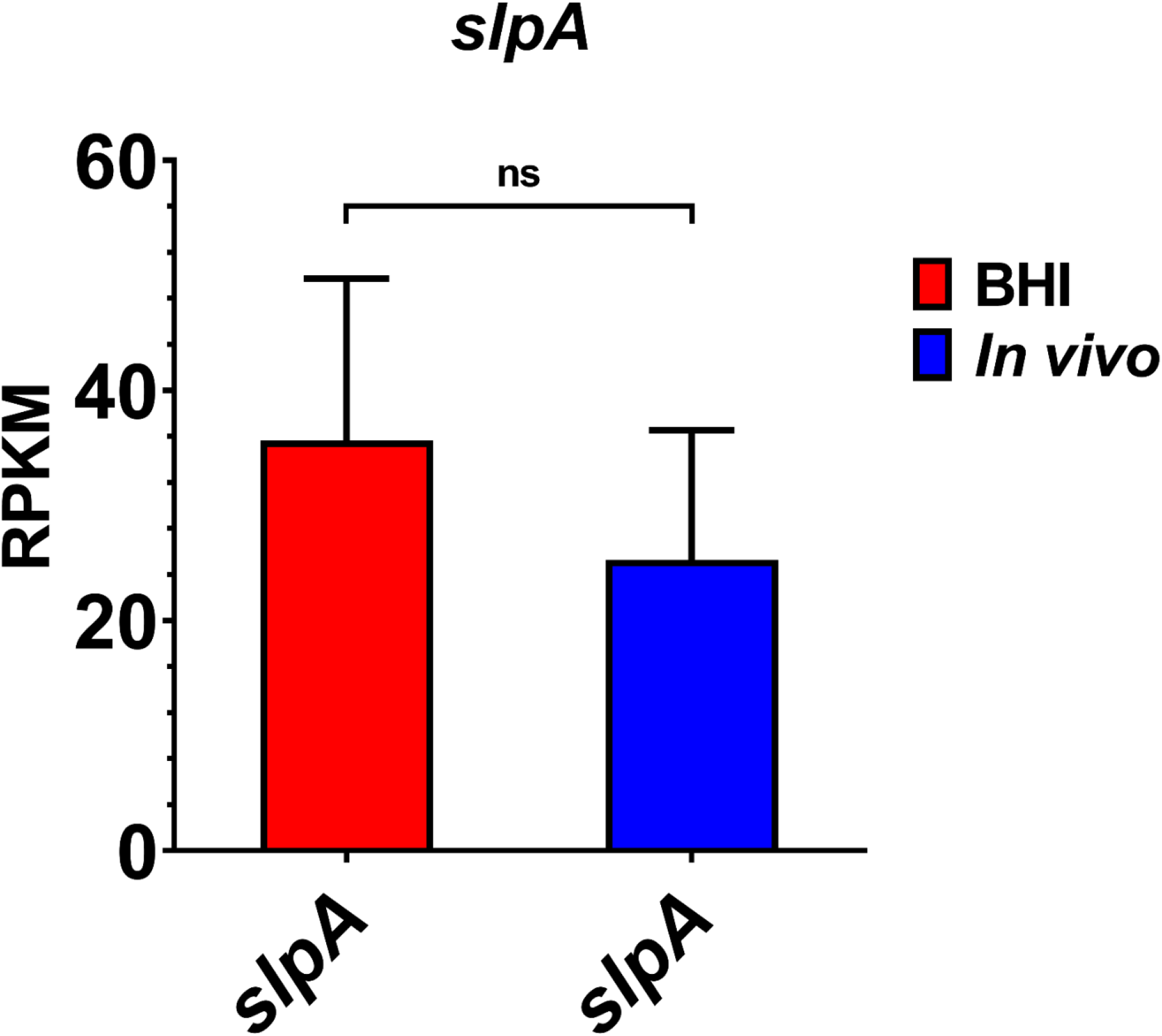
Normalized surface layer protein (SlpA) gene expression in BHI or *in vivo.* Reads per kilobase per million (RPKM) for the *slpA* gene (ns = not significant) *in vivo* relative to BHI. BHI is shown in red, and *in vivo* in blue. RPKM values are the means ± the standard deviations of three independent RNA-Seq runs.

## Discussion

The objective of the current study was to identify known and putative *B. cereus* virulence factor expression in a mouse model of endophthalmitis, and compare this to expression in laboratory medium BHI. Eight hours postinfection was evaluated because we have consistently observed significant inflammatory cell influx and retinal architectural damage at this time point. In both rabbit and mouse endophthalmitis models, *B. cereus* replication upon entry into the vitreous and dissemination throughout the globe occurs rapidly [12, 17, 23]. As early as 4 hours postinfection, inflammatory cells (primarily neutrophils) enter the eye but fail to control the growth of *B. cereus.* At 12 to 18 hours following bacterial entry, a complete dissolution of the retinal architecture and retinal function loss is observed [23]. *B. cereus* endophthalmitis in human patients also occurs following a similarly rapid course, and results in loss of vision or complete loss of the globe in 12 to 48 hours [2–5]. Evaluation of the virulome of *B. cereus in vivo* might lead to the identification of *B. cereus* factors that contribute to immune cell interference and pathogenesis.

Hbl and Nhe are both pore-forming toxins regulated by the PlcR/PapR system, and have known roles in *B. cereus* gastrointestinal infections. Both sets of toxin genes were expressed at 8 hours postinfection in our murine model of *B. cereus* endophthalmitis. Two lytic components, L1 (*hblD*) and L2 *(hblC),* and a single binding component, B *(hblA)* comprise the secreted, hemolytic Hbl toxin [26, 27]. Hbl toxin expression was observed among subsets of isolates that caused gastrointestinal infections [28, 29], and the *hblCA* genes were detected in endophthalmitis and keratitis isolates by PCR [30]. In a rabbit model of *B. cereus* endophthalmitis, no difference in infection progression was observed between a wild type *B. cereus* and an isogenic Hbl-deficient mutant [11]. However, injection of purified Hbl into rabbit eyes resulted in pathological effects similar to those observed during infection with live bacteria [27]. Taken together, these results suggested that additional virulence factors might contribute to retinal damage and the unique inflammatory response to *B. cereus* infection. Here, Hbl expression was significantly higher at 8 hours postinfection than in BHI. We cannot rule out that cytotoxicity *in vivo* might be attributable to a combination of *B. cereus* secreted toxins. Nhe is also a tripartite toxin consisting of the lytic components NheA *(nheL2)* and NheB (*nheL1*), and the binding component NheC [31]. Nhe was shown to be expressed in the majority of gastrointestinal-associated isolates [28, 29]. We previously observed lower level expression of these toxins in *ex vivo* vitreous than in laboratory media [20]. In the current analysis, Hbl and Nhe were expressed at significantly higher levels at 8 hours postinfection, suggesting that the *in vivo* environment influences expression of these genes.

We also evaluated the expression of the putative cell wall peptidase-encoding gene *entFM*, and two putative enterotoxin/cell-wall binding protein genes *entA*, and *entC* at the midpoint of endophthalmitis progression. Similar levels of the *entFM* and *entC* genes were detected *in vivo* and after 8 hours of growth in BHI. However, expression of *entA* was higher at the midpoint of infection than after growth in BHI. EntFM contributed to vacuolization in macrophages, *in vitro* adhesion and biofilm formation, and lethality in a *Galleria mellonella* model [32], it is not clear whether EntA or EntC contribute to virulence in any model.

Expression of the HlyA-encoding gene *hlyA* was similar *in vivo* and in BHI.However the cereolysin O-encoding gene *cerO* was expressed at significantly higher levels *in vivo* than in BHI. Cereolysin O (CerO) is a member of the cholesterol-dependent cytolysin (CDC) family of pore-forming toxins. At 4 hours postinfection with a CerO-deficient *B. cereus* strain, TLR4-dependent inflammatory mediators were significantly downregulated [33]. TLR4 contributed to the inflammatory response to *B. cereus* [22], and was activated by purified *B. cereus* SlpA protein [18]. In addition to direct cellular cytotoxicity, CerO as a CDC might interact with and/or activate TLR4 [34–37], inciting inflammation during *B. cereus* endophthalmitis.

*In vivo,* the metalloprotease genes *inhA1* and *inhA2* genes were expressed at similar levels. Expression of *inhA1* and *inhA2* were also similar to one another in *ex vivo* vitreous [20]. *InhA1* was expressed at significantly higher levels *in vivo* than in BHI, however, *inhA2* was expressed at similar levels in both environments. We did not observe expression of *inhA3* in either environment, similar to our previous study that did not detect expression in BHI, LB, or *ex vivo* vitreous [20]. These observations concur with the report that *B. cereus* ATCC 14579 does not express *inhA3* due to a mutation in *npr* [25]. The cell surface-associated metalloproteinase, camelysin (CalY), was also not expressed in either environment, in contrast to our previous study that expression was detected in LB and *ex vivo* vitreous [20]. The InhA metalloproteases have been proposed to contribute to virulence by mediating degradation of extracellular matrix components and escaping host defenses [38]. Based on these results, it might be postulated that the InhA metalloproteases might damage the eye by breaking down type II collagen in the vitreous [39], or by mediating escape from neutrophils. CalY degrades serum protease inhibitors, collagen type I, fibrin, fibrinogen, and plasminogen [40, 41]. While we reported that *calY* expression was similar to *inhA* expression after growth in the vitreous environment, *calY* transcripts were not detected at 8 hours postinfection. However, these results do not preclude the involvement of CalY in pathogenesis, as expression might occur at a different time point during intraocular infection.

Among the transcriptional regulators analyzed, only *codY*, *gntR*, and *nprR* were expressed at 8 hours postinfection. *gntR* and *nprR* were the most highly expressed. CodY is a global transcriptional regulator that affects both the PlcR/PapR and Spo0A-AbrB regulatory systems [42]. Deletion of *codY* decreased the expression of PlcR-regulated virulence genes, resulting in attenuated infection in a *G. mellonella* model [42]. On the contrary, deletion of *codY* in *Staphylococcus aureus* did not alter its intraocular growth but did result in significantly higher inflammatory scores and lower retinal function retention as compared to the parental wild type strain [43]. We reported that *codY* was expressed in *ex vivo* vitreous as well as in BHI and LB media during stationary phase [20]. In the current study, we detected *codY* transcripts at the midpoint of endophthalmitis. Given that CodY binds to the promoter of InhA1 [42], our results suggest that CodY might play an integral role in the *in vivo* regulation of *inhA1.* Moreover, CodY might activate the expression of secreted proteases involved in extracellular matrix degradation and thus promote physical barrier breach in the eye. CodY might also regulate the expression of metabolic pathway genes that are needed to utilize available nutrient sources during intraocular infection.

The gluconate transcriptional repressor, GntR, was expressed *in vivo* at the 8-hour time point. Expression was also observed after 18 hours of growth in BHI, LB, and *ex vivo* vitreous [20]. GntR represses transcription of the gluconate operon *(gntRKPZ)* which encodes proteins necessary to utilize gluconate as a carbon source, and repression is relieved by the presence of gluconate [44]. The diarrheal pathogens *Vibrio cholera* and *Escherichia coli* are able to adapt to the gastrointestinal tract by utilization of available gluconate [45]. In the absence of gluconate, intraocular expression of *gntR* would be expected to repress the expression of the other three genes necessary to metabolize gluconate.

The quorum sensing transcriptional regulator gene, *nprR*, was expressed at significantly higher levels at 8 hours postinfection than at the same time point in BHI. *In vitro* analysis in our prior study detected *nprR* transcripts during in stationary phase in BHI, LB, and *ex vivo* vitreous. At the start of stationary phase, NprR regulates PlcR as well as genes involved in cell survival, sporulation, antibiotic resistance, and the metalloprotease *nprA* gene [46]. In strain ATCC 14579, *nprR* is disrupted by a transposon insertion, resulting in a nonfunctional polypeptide. Dubois et al. reported that NprR positively regulates the *inhA3* expression [25]. Our results suggest that the absence of *inhA3* transcripts i*n vitro* and *in vivo* was likely due to a nonfunctional NprR in ATCC 14579. The *nprR* gene is also repressed by CodY during logarithmic phase growth, an observation consistent with detecting low-level expression at 8 hours in BHI. However, in eye, *nprR* expression was 56-fold higher than in BHI, suggesting that expression of *nprR* is influenced by *in vivo* queues.

In conjunction with PlcR, SinR regulates the constitutive expression of Hbl in *B. thuringensis* biofilms [47]. However, while *hbl* gene expression was high at the midpoint of infection in our murine model, *sinR* transcripts were absent. This is in contrast to high levels in BHI at 8 hours and after growth to stationary phase in BHI, LB, or *ex vivo* vitreous [20]. We previously observed low levels of expression of *sinI,* the gene encoding the inhibitor of SinR [48], after growth to stationary phase in BHI and *ex vivo* vitreous, but not after growth to stationary phase in LB [20]. Similarly, *sinI* transcripts were not detected *in vivo* or during logarithmic phase growth in BHI. The absence of *sinR* transcripts at 8 hours postinfection again suggests that *in vivo*, *hbl* expression might be governed by other factors.

Transcripts of *entD,* the gene encoding the EntD biofilm regulator, were detected at low levels during stationary phase after 18 hours of growth in BHI, LB, and *ex vivo* vitreous [20], but were not detected at 8 hours in BHI or following intraocular infection. EntD is involved in regulating genes involved in biofilm formation, cell metabolism, cell structure, antioxidation, motility, and toxin production [49]. This suggests that EntD might contribute to intraocular survival. However, given the differences between our previous and current findings, low level expression of *entD* might only occur at later stages of infection and thus the relevance to intraocular infection remains unknown.

The PlcR/PapR master quorum sensing regulatory system transactivates a subset of virulence factors at the beginning of stationary phase. Low nutrient conditions and increasing cellular density result in activation of *plcR* expression [31, 50]. Surprisingly, we did not detect transcripts of either the *plcR* or the *papR* genes *in vivo* at 8 hours postinfection. In conjunction with the rapid growth of *B. cereus* that occurs upon infection of the eye, these results suggest that nutrients are either not limiting, and/or sufficient cell densities have not yet been reached at this time point. *plcR* expression was observed at the end of the exponential growth phase in LB [51], and given that we previously detected low levels of transcripts in stationary phase after growth in BHI, LB, and *ex vivo* vitreous [20], expression of *plcR in vivo* might occur during narrow window at the end of logarithmic and beginning of stationary phase. Further, *plcR* expression in the eye during infection is supported by our report that PlcR contributes to inflammation and retinal function loss during experimental *B. cereus* endophthalmitis [9], and permeability of an *in vitro* blood retinal barrier [52]. Our *in vivo* results at this timepoint also suggests that virulence factors controlled by PlcR *in vitro* might be regulated differently, or respond to different environmental queues during infection.

*B. cereus* motility is important to its virulence during endophthalmitis [8, 19]. We therefore evaluated expression of genes related to motility and chemotaxis at the midpoint of endophthalmitis. These genes reside in 45 kb region in the *B. cereus* ATCC14579 genome [53, 54], and, in *B. subtilis,* are organized in a single operon under the control of CodY [55, 56]. During stationary phase in *ex vivo* vitreous, the chemotaxis-related genes *cheA, cheY,* and *cheR,* were expressed at low levels, and expression of the genes related to flagellum production and motility *fla, fliF,* and *motB* was not detected or detected at only at low levels [20]. In our current study, we detected low levels of *cheA* transcripts, but did not detect *cheY* or *cheR* transcripts at 8 hours following infection. However, the motility-related genes *motB*, *fliF*, and *fla* expression was higher at this timepoint than after 18 hours of growth in *ex vivo* vitreous [20]. CheA, CheY, and CheR are involved in switching between tumbling and forward motion in order to adjust to chemoattractant gradients [53, 57–60]. In the eye, nutrients might be homogeneously distributed throughout the vitreous, so following a chemoattractant gradient is not necessary. However, detection of motility-related gene expression at the midpoint of infection supports our previous studies demonstrating the importance of motility to endophthalmitis pathogenesis [8, 19]. These findings suggest that cues necessary for expression of motility genes in the eye were absent from *ex vivo* vitreous [20].

Similar to our findings in *ex vivo* vitreous [20], the *sodA2* gene was the most highly expressed among the genes we evaluated *in vivo* at 8 hours postinfection. SodA2 is a Mn-based superoxide dismutase [61] that mediates the conversion of superoxide anion (O2^-^) and other oxygen radicals to molecular oxygen and hydrogen peroxide (H2O2). These products are then converted into H2O and O2 by peroxidases and catalases. *B. subtilis* SodA provides protection from oxidative stress in growing and sporulating cells [62]. Bacterial SodA might serve as a novel therapeutic target given that human SOD is structurally different [63]. High level expression of the *B. cereus sodA* genes, in particular *sodA2,* in the eye during infection might interfere with neutrophil function and impede clearance. Indeed, *Bacillus* replicate seemingly without much opposition, even as neutrophils are recruited rapidly and to high numbers into the eye during infection.

An additional target for therapeutic intervention in *B. cereus* endophthalmitis is the surface layer protein, SlpA. We identified SlpA as a significant contributor to disease severity in our murine model of *Bacillus* endophthalmitis [17]. The mechanism for this effect was linked to activation of nuclear factor kappa-light-chain-enhancer of activated B cells (NF-κB) and subsequent inflammatory mediator production from Muller cells [17]. Further investigation revealed that SlpA activated both TLR2 and TLR4 *in vitro,* and that administration of TLR2/4 inhibitors attenuated the severity of infection in our mouse model (18, 24). These results suggested that interfering with SlpA-mediated innate immune activation might serve as a novel target for immunomodulatory treatments of *B. cereus* endophthalmitis. Although *slpA* transcript levels were not significantly different after 8 hours of growth *in vivo* or in BHI, detection of transcripts *in vivo* does support our findings that S-layer protein is present and contributes to the host immune response and poor vision outcomes in a mouse model of *Bacillus* endophthalmitis.

In summary, our results identified the expression of virulence genes and virulence-associated genes at the midpoint of *Bacillus* infection in the eye, highlighting possible mechanisms for inflammation, retinal damage, and vision deficits in this disease. While our results provide a snapshot of steady state transcript levels at a single, but clinically important time point during infection, these findings lay the groundwork for and demonstrate the feasibility of examining the dynamic *B. cereus* virulome during the course of endophthalmitis, and further analysis of potential targets for anti-virulence therapies to combat this devastating infection.

## Abbreviations

BHI: Brain Heart Infusion;
RNA-Seq: RNA Sequencing;
POE: post-operative endophthalmitis;
PTE: post-traumatic endophthalmitis;
EE: endogenous endophthalmitis;
PC-PLC: phosphatidylcholine-specific phospholipase C;
PI-PLC: phosphatidylinositol-specific phospholipase C;
TLR: Toll-like receptor;
Hbl: hemolysin BL;
Nhe: nonhemolytic enterotoxin;
Ent: enterotoxin;
InhA: immune inhibitor A;
CerO: cereolysin O;
HylA: hemolysin A;
SOD: superoxide dismutase;
Slp: surface layer protein;
RPKM: reads per kilobase million;
bp: basepairs.

## Author Statements

### Authors and contributors

PSC, FCM, and MCC designed the study. PSC, FCM, MAE, CL, ALL, and MHM performed the experiments. PSC, FCM, MAE, and CL compiled and analyzed the data. PSC performed the statistical analyses on the data. PSC prepared the original draft of the manuscript. FCM and MCC reviewed and edited the manuscript. PSC and MCC managed the project. MCC acquired the funding for this project. All authors read and approved the final manuscript.

## Acknowledgements

We thank Roger Astley (Department of Ophthalmology, OUHSC) and Mark Dittmar (Dean McGee Eye Institute Animal Facility) for their invaluable technical assistance. We thank Jenny Gipson and Allison Gillaspy at the Laboratory for Molecular Biology and Cytometry Research at OUHSC for assistance and the use of the Core Facility which provided the RNA-Seq service.

